# Single-cell transcriptome analysis reveals mesenchymal stem cells in cavernous hemangioma

**DOI:** 10.1101/2021.09.02.458742

**Authors:** Fulong Ji, Yong Liu, Jinsong Shi, Chunxiang Liu, Siqi Fu, Heng Wang, Bingbing Ren, Dong Mi, Shan Gao, Daqing Sun

**Affiliations:** Department of Paediatric Surgery, Tianjin Medical University General Hospital, Tianjin 300052, P.R. China; College of Life Sciences, Nankai University, Tianjin, Tianjin 300071, P.R. China; National Clinical Research Center of Kidney Disease, Jinling Hospital, Nanjing University School of Medicine, Nanjing, Jiangsu 210016, P.R. China; School of Mathematical Sciences, Nankai University, Tianjin, Tianjin 300071, P.R. China

**Keywords:** MSC, Vascular malformation, Vascular tumor, Stem cell, scRNA-seq

## Abstract

A cavernous hemangioma, well-known as vascular malformation, is present at birth, grows proportionately with the child, and does not undergo regression. Although a cavernous hemangioma has well-defined histopathological characteristics, its origin and formation remain unknown. In the present study, we characterized the cellular heterogeneity of cavernous hemangioma using single-cell RNA sequencing (scRNA-seq). The main contribution of this study is the discovery of mesenchymal stem cells (MSCs) that cause tumour formation in cavernous hemangioma and we propose that these MSCs may be abnormally differentiated or incompletely differentiated from epiblast stem cells.

Other new findings include the responsive ACKR1 positive endothelial cell (ACKR1+EC) and BTNL9 positive endothelial cell (BTNL9+EC) and the BTNL9-caused checkpoint blockade enhanced by the *CXCL12-CXCR4* signalling. The activated CD8+T and NK cells may highly express *CCL5* for their infiltration in cavernous hemangiomas, independent on the tumor cell-derived *CCL5-IFNG*-*CXCL9* pathway. The highly co-expression of *CXCR4* and *GZMB* suggested that plasmacytoid dendritic cells (pDCs) function for anti-tumour as CD8+T cells in cavernous hemangiomas. The oxidised low-density lipoprotein (oxLDL) in the TME of cavernous hemangiomas may play an important role as a signalling molecular in the immune responses. Notably, we propose that oxLDL induces the oxLDL-OLR1-NLRP3 pathway by over-expression of *OLR1* in M1-like macrophages, whereas oxLDL induces the oxLDL-SRs-C1q (SRs are genes encoding scavenger receptors of oxLDL except *OLR1*) pathway by over-expression of other scavenger receptors in M2-like macrophages.

The present study revealed the origin of cavernous hemangiomas and discovered marker genes, cell types and molecular mechanisms associated with the origin, formation, progression, diagnosis or therapy of cavernous hemangiomas. The information from the present study makes important contributions to the understanding of cavernous hemangioma formation and progression and facilitates the development of gold standard for molecular diagnosis and effective drugs for treatment.

## Introduction

Vascular tumours include hemangioma, hemangioendothelioma, angiosarcoma, and their epithelioid variants [1]. According to the size of the affected vessels, hemangiomas are histologically classified as capillary, cavernous, or mixed-type hemangiomas [2]. A capillary hemangioma (superficial, red, raised), also called strawberry hemangioma, is a tumour of infancy that undergoes a phase of rapid growth and expansion followed by a period of slow but steady regression during childhood. In contrast, a cavernous hemangioma (deep dermal, blue hue), which is now classified as vascular malformation according to the International Society for the Study of Vascular Anomalies (ISSVA) classification [3], is present at birth, grows proportionately with the child, and does not undergo regression [4]. Major features used to discriminate cavernous hemangiomas from capillary hemangiomas include the observation of “normal” vascular endothelial cells in cavernous hemangiomas and the over-expression of vascular endothelial growth factor A (*VEGFA*) and fibroblast growth factor receptor 1 (*FGFR1*) in capillary hemangiomas during the proliferative stage [6]. Cavernous hemangiomas have been reported to arise at various sites, including the skin and subcutaneous layers of the head and neck, face, extremities, liver, gastrointestinal tract, and even the thymus [5]. The tumours are composed of dilated vascular spaces, with thinned smooth muscle walls separated by a variable amount of fibroconnective tissue.

Three classes of cavernous hemangiomas hepatic cavernous hemangioma (HCH), retinal cavernous hemangioma (RCH), and cerebral cavernous hemangioma (CCH), also known as cerebral cavernous malformation (CCM), are comparatively well studied. HCH, the most common benign tumour of the liver, is present in up to 7% of individuals that participate in autopsy studies. Histological examination of the lesions revealed a network of vascular spaces lined by endothelial cells and separated by a thin fibrous stroma. Large HCHs may be associated with thrombosis, scarring, and calcification [7]. RCH is composed of clusters of saccular aneurysms filled with dark blood. Microscopic examination of the lesions revealed multiple thin-walled interconnected vascular spaces lined by flat endothelial cells, with red cell necrosis and partially organised intravascular thrombosis. Vascular spaces are bordered by thin, fibrous septa, with occasional nerve fibers and glial cells. Although mutations in *KRIT1* and *CCM1* genes have been found in patients with both RCH and CCH [8], the cause of RCH remains unknown. CCHs that occur in the central nervous system, most often in the brain, can cause intracranial hemorrhage, seizures, neurological deficits, and even death. Ultrastructural studies revealed abnormal or absent blood–brain barrier components, poorly formed tight junctions with gaps between endothelial cells, lack of astrocytic foot processes, and few pericytes in the lesions. CCH has sporadic and familial forms; familial CCHs often display multiple lesions and autosomal dominant inheritance. According to the current theory, capillary hemangiomas originate from neogenesis or revival of dormant embryonic angioblasts and arise through hormonally driven vessel growth [4], while familial CCHs may be caused by loss-of-function mutations in three genes, *KRIT1, CCM2*, and *PDCD10* [9]. However, the origins of capillary and cavernous hemangiomas remain controversial and unknown, respectively.

Although capillary hemangiomas are mainly treated via surgery, several drugs (e.g., propranolol and glucocorticoids) have been developed to avoid the risks of intraoperative profuse bleeding, postoperative recurrence, long-term scarring, and other complications. The treatment of cavernous hemangiomas is still dependent on excision and venous embolization therapy [3]; however, inadequate excision causes recurrence of cavernous hemangiomas [10]. To develop drugs or other new therapies, more basic research must be conducted to understand cavernous hemangiomas at the molecular level. In the present study, we characterised the cellular heterogeneity of cavernous hemangioma using single-cell RNA sequencing (scRNA-seq) [11]. Through further analysis of the scRNA-seq data, we aimed to: (1) reveal the comprehensive cellular composition and gene expression profile of a cavernous hemangioma at the single-cell level, and (2) discover marker genes, cell types, and molecular mechanisms associated with the origin, formation, progression, diagnosis, and therapy of cavernous hemangiomas.

## Results

### Single-cell RNA sequencing and basic analyses

The lesion was obtained as tumour tissue from a 6-year-old patient diagnosed with cavernous hemangioma (**Supplementary File 1**). Using tissue in the center of the lesion, scRNA-seq libraries (10x Genomics, USA) were constructed and sequenced to produce ∼163 Gbp of raw data (**Materials and Methods**). After data cleaning and quality control, a total of 10,784 cells and 22,021 genes were identified to produce a 22,010×10,784 matrix and an 11×10,784 matrix, representing the expression levels of nuclear and mitochondrial genes, respectively. Using the 22,010×10,784 nuclear matrix, 10,784 cells were clustered into 18 clusters with adjusted parameters (**Materials and Methods**) and then merged into 16 clusters (**Figure 1A**), including fibroblast cell type 1 (fibroblast1), type 2 (fibroblast2), smooth muscle cell (SMC), endothelial cell type 1 (EC1), type 2 (EC2), lymphatic endothelial cell (LEC), T lymphocyte type 1 (TC1), type 2 (TC2), type 3 (TC3), B lymphocyte (BC), mast cell, monocyte derived dendritic cell (mDC), plasmacytoid dendritic cell (pDC), CLEC9A positive dendritic cell (CLEC9A^+^DC), macrophage type 1 (Maph1) and type 2 (Maph2), respectively. By using known marker genes, 16 clusters were identified as 16 main cell types and then renamed as fibroblast, mesenchymal stem cell (MSC), SMC, EC1, EC2, LEC, CD4 positive T cell (CD4+TC), CD8 positive T cell (CD8+TC), natural killer cell (NKC), BC, mast, mDC, pDC, CLEC9A^+^DC, m1Maph and m2Maph clusters, respectively. To confirm the presence of the main cell types, dilated capillaries, normal capillaries, lymphatic vessels, muscle tissue, and connective tissue were observed by haematoxylin–eosin (HE) staining (**Supplementary file 1**).

**Figure 1.**
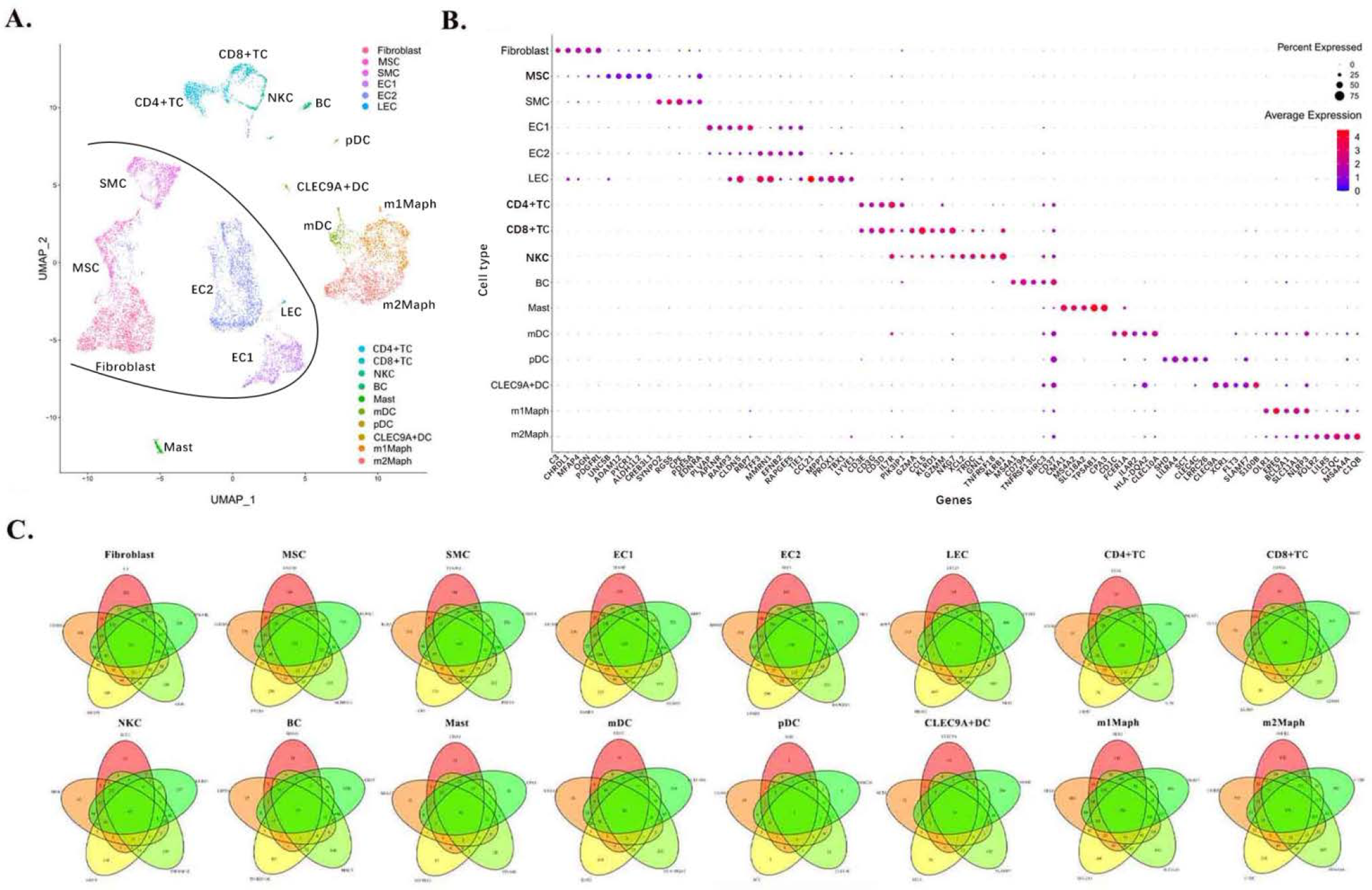
Identification of main cell types in cavernous hemangioma. A total of 10,784 cells were clustered into 16 clusters (fibroblast, MSC, SMC, EC1, EC2, LEC, CD4+TC, CD8+TC, NKC, BC, mast, mDC, pDC, CLEC9A+ DC, m1Maph and m2Maph) that were identified as 16 main cell types. **A**. Uniform Manifold Approximation and Projection (UMAP) method was used to show the clustering results.; **B**. For each of 16 clusters, a combination of five marker genes (**Table 1**) were assigned. The differential expression analysis between the cells inside and outside the cluster was performed to select top five differentially expressed (DE) genes as the marker genes; **C**. The representation of a cell type by the combination of marker genes were also showed in Venn diagrams. MSC: mesenchymal stem cell, SMC: smooth muscle cell, EC1: endothelial cell type 1, EC2: endothelial cell type 2, LEC: lymphatic endothelial cell, CD4+TC: CD4 positive T cell, CD8+TC: CD8 positive T cell, NKC: natural killer cell, BC: B cell, mDC: monocyte derived dendritic cell, pDC: plasmacytoid dendritic cell, CLEC9A+DC: CLEC9A positive dendritic cell, m1Maph: M1-like macrophage and m2Maph: M2-like macrophage.

For each of the 16 clusters, the cell type was identified by comparing the selected differentially expressed (DE) genes to known marker genes. Differential expression analysis between cells inside and outside the cluster (**Supplementary file 2**) was performed to select DE genes based on the ratio between the percentage of cells expressing the gene inside (PCTin) and outside the cluster (PCTout) (**Materials and Methods**). As no single marker gene can be used to discriminate a cell type from its relatives (e.g., CD8+TC from CD4+TC), we used a combination of five marker genes (**Table 1**) to improve the identification of cell types. By sorting the ratio between PCTin and PCTout in descending order, the top five DE genes were selected as a combination of marker genes for each cell type (**Figure 1B**). The representation of a cell type by the combination of marker genes was also presented in Venn diagrams (**Figure 1C**). We designed a new metric (**Materials and Methods**), called the union and intersection coverage of a cluster (UICC), to evaluate the representation of a cell type by a combination of marker genes (**Table 1**).

**Table 1.**
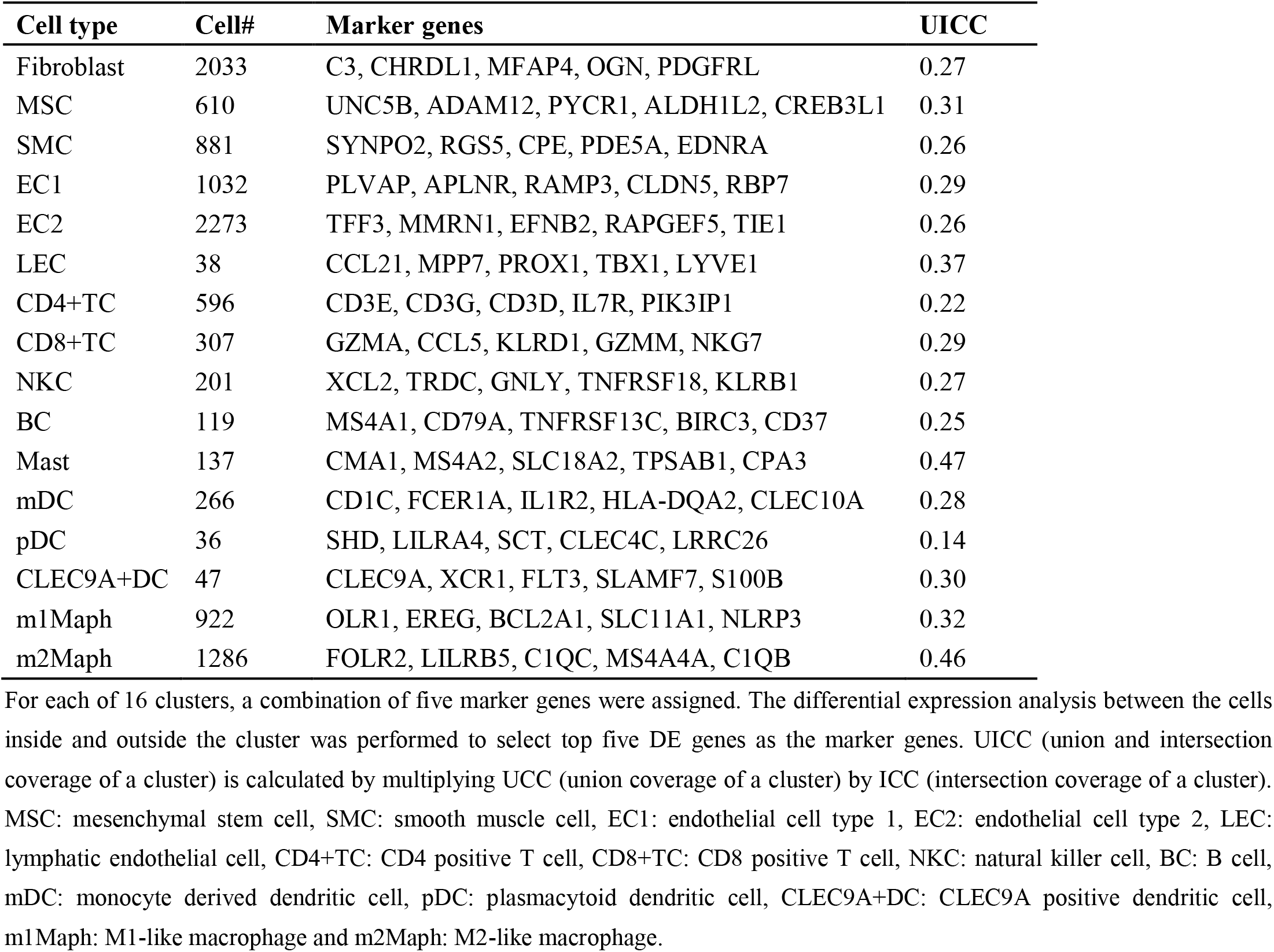
Marker genes of 16 main cell types.

| Cell type | Cell# | Marker genes | UICC |
| --- | --- | --- | --- |
| Fibroblast | 2033 | C3, CHRDL1, MFAP4, OGN, PDGFRL | 0.27 |
| MSC | 610 | UNC5B, ADAM12, PYCR1, ALDH1L2, CREB3L1 | 0.31 |
| SMC | 881 | SYNPO2, RGS5, CPE, PDE5A, EDNRA | 0.26 |
| EC1 | 1032 | PLVAP, APLNR, RAMP3, CLDN5, RBP7 | 0.29 |
| EC2 | 2273 | TFF3, MMRN1, EFNB2, RAPGEF5, TIE1 | 0.26 |
| LEC | 38 | CCL21, MPP7, PROX1, TBX1, LYVE1 | 0.37 |
| CD4+TC | 596 | CD3E, CD3G, CD3D, IL7R, PIK3IP1 | 0.22 |
| CD8+TC | 307 | GZMA, CCL5, KLRD1, GZMM, NKG7 | 0.29 |
| NKC | 201 | XCL2, TRDC, GNLY, TNFRSF18, KLRB1 | 0.27 |
| BC | 119 | MS4A1, CD79A, TNFRSF13C, BIRC3, CD37 | 0.25 |
| Mast | 137 | CMA1, MS4A2, SLC18A2, TPSAB1, CPA3 | 0.47 |
| mDC | 266 | CD1C, FCER1A, IL1R2, HLA-DQA2, CLEC10A | 0.28 |
| pDC | 36 | SHD, LILRA4, SCT, CLEC4C, LRRC26 | 0.14 |
| CLEC9A+DC | 47 | CLEC9A, XCR1, FLT3, SLAMF7, S100B | 0.30 |
| m1Maph | 922 | OLR1, EREG, BCL2A1, SLC11A1, NLRP3 | 0.32 |
| m2Maph | 1286 | FOLR2, LILRB5, C1QC, MS4A4A, C1QB | 0.46 |

For each of 16 clusters, a combination of five marker genes were assigned. The differential expression analysis between the cells inside and outside the cluster was performed to select top five DE genes as the marker genes. UICC (union and intersection coverage of a cluster) is calculated by multiplying UCC (union coverage of a cluster) by ICC (intersection coverage of a cluster). MSC: mesenchymal stem cell, SMC: smooth muscle cell, EC1: endothelial cell type 1, EC2: endothelial cell type 2, LEC: lymphatic endothelial cell, CD4+TC: CD4 positive T cell, CD8+TC: CD8 positive T cell, NKC: natural killer cell, BC: B cell, mDC: monocyte derived dendritic cell, pDC: plasmacytoid dendritic cell, CLEC9A+DC: CLEC9A positive dendritic cell, m1Maph: M1-like macrophage and m2Maph: M2-like macrophage.

In a previous study of ascending thoracic aortic aneurysms (ATAAs) [12], 11 major cell types were identified by known marker genes, including fibroblast, MSC, SMC1, SMC2, EC, TC, NKC, BC, mast cell, MonoMaphDC (monocyte/macrophage/DC) and plasma cell. A simple comparison between cell populations in cavernous hemangioma tissue and those in ATAA tissue and their controls [12] revealed several main differences: (1) cavernous hemangioma tissue contained fewer immune cells (36.32% of the total cells) than ATAA tissue (64.08%) and more than the controls of ATAA tissue (26.24%); (2) cavernous hemangioma tissue contained significantly higher proportions of ECs and fibroblasts (30.65% and 24.51%) than ATAA tissues (7.43% and 7.6%) and their controls (14.02% and 13.51%); (3) cavernous hemangioma and ATAA tissue contained more macrophages and DC cells (23.71% and 21.73% of the total cells) than the ATAA tissue controls (7.64%); and (4) ATAA tissue contained significantly more B and plasma cells than their controls and cavernous hemangioma tissue.

### Discovery of MSCs

The fibroblast1 and fibroblast2 clusters contained 18.85% (2,033/10,784) and 5.66% (610/10,784) of the total cells, respectively. Such proportion of fibroblasts was markedly higher than that of SMCs (8.17%, 881/10784), which is consistent with the prominent histological features of cavernous hemangiomas where the thinned smooth muscle walls are separated by a variable amount of fibroconnective tissue (**Introduction**). Although both fibroblast1 and fibroblast2 expressed a number of fibrillin, fibulin, collagen, and elastin genes required in the extracellular matrix (ECM), including *FBN1, FBLN1, FBLN2, FBLN5, COL1A1, COL1A2, COL3A1, COL6A1, COL6A2, COL5A2, COL14A1*, and *ELN* (**Figure S4**), fibroblast1 showed significantly higher expression of most of these genes (e.g., *FBLN1, FBLN2*, and *ELN*) compared with fibroblast2. Fibroblast1 also highly expressed all the fibroblast marker genes, *PDGFRA, PDGFRB, MEG3, SCARA5, COL14A1*, and *OGN* (**Figure S5**), which was reported in a previous study [13]. In contrast, fibroblast2 expressed *PDGFRA, PDGFRB, MEG3*, and *COL14A1* at low levels and *SCARA5* and *OGN* at very low levels. According to the previous study [13], the marker genes of early (*MEG3* and *SCARA5*), intermediate (*COL14A1* and *OGN*), and terminal states via pericyte-to-myofibroblast differentiation (*NKD2, GREM2, NRP3*, and *FRZB*) are highly expressed in myofibroblasts. These results indicate that fibroblast1 is a cluster of fibroblasts, whereas fibroblast2 belongs to cell types related to fibroblasts. Both the fibroblast1 and fibroblast2 clusters did not originate from pericytes, as *NKD2, GREM2*, and *FRZB* were barely detected and *NRP3* was not detected (**Figure S5**).

Differential expression analysis between cells inside and outside fibroblast2 was performed to generate a gene-expression signature (**Supplementary file 2**), which merits further studies, particularly on the development of potential targets for the diagnosis or treatment of cavernous hemangiomas. Using LFCio above 2 (**Materials and Methods**), 63 coding genes and a long noncoding RNA (lncRNA) gene, *RP11-14N7*.*2* (**Table 2**), were selected for further analysis. Gene ontology (GO) and pathway annotation of 63 coding genes (**Figure 2A**) revealed that 57.14% (36/63) of the genes were involved in ECM organisation (GO:0030198), and 15.87% (10/63) of the genes were involved in the response to fibroblast growth (GO:0071774). This finding confirmed that fibroblast2 is a cell type related to fibroblasts. Importantly, many GO annotations were found to be enriched in endodermal cell differentiation and tissue development, including endodermal cell differentiation (GO:0035987), blood vessel development (GO:0001568), skeletal system development (GO:0001501), heart development (GO:0007507), bone development (GO:0060348), muscle organ development (GO:0007517), reproductive structure development (GO:0048608), and lung development (GO:0030324). In particular, *TWIST1, SULF1, COL5A1, COL5A2, COL6A1*, and *FN1* were associated with endodermal cell differentiation (GO:0035987), thus indicating that fibroblast2 exhibits characteristics of stem or progenitor cells. A further investigation of the 63 coding genes (**Table 2**) revealed that at least 19 genes were over-expressed or up-regulated in stem cells, namely, *UNC5B, ADAM12, PYCR1, PHGDH, SLIT2, PFN2, TNFAIP6, TNC, EDIL3, TWIST1, SNAI2, PHLDA2, LOXL2, BMP1, COL5A1, POSTN, ID3, COL6A1*, and *COL1A1* (**Figure 2B**). Among these genes, *PYCR1, TNFAIP6, EDIL3, TWIST1, LOXL2, BMP1*, and *COL1A1* were reported to be expressed in MSCs according to previous studies (**Supplementary file 3**). *TWIST1* is a basic helix-loop-helix (bHLH) transcription factor that plays essential and pivotal roles in multiple stages of embryonic development. The over-expression of *TWIST1* induces epithelial-mesenchymal transition (EMT), a key process in cancer metastasis [14]. Among the five marker genes (**Table 1**), *UNC5B* was reported to be a marker gene of epiblast stem cells. According to annotations from the GeneCards database [15], *UNC5B* encodes a member of the netrin family of receptors, and the encoded protein mediates the repulsive effect of netrin-1. The protein encoded by *UNC5B* belongs to a group of proteins called dependence receptors (DpRs), which are involved in embryogenesis and cancer progression. These findings indicate that fibroblast2 is a cluster of MSCs that may be abnormally differentiated or incompletely differentiated from epiblast stem cells and cause cavernous hemangioma. Fibroblast2 is not a cluster of cancer-associated fibroblasts (CAFs), as the CAF marker genes *VEGFA, VEGFB, VEGFC, HGF, GAS6, TGFB2, TGFB3, IL6*, and *CXCL12* [16] were barely detected, and *TGFB1* was expressed at the medium level in the fibroblast2 cluster.

**Figure 2.**
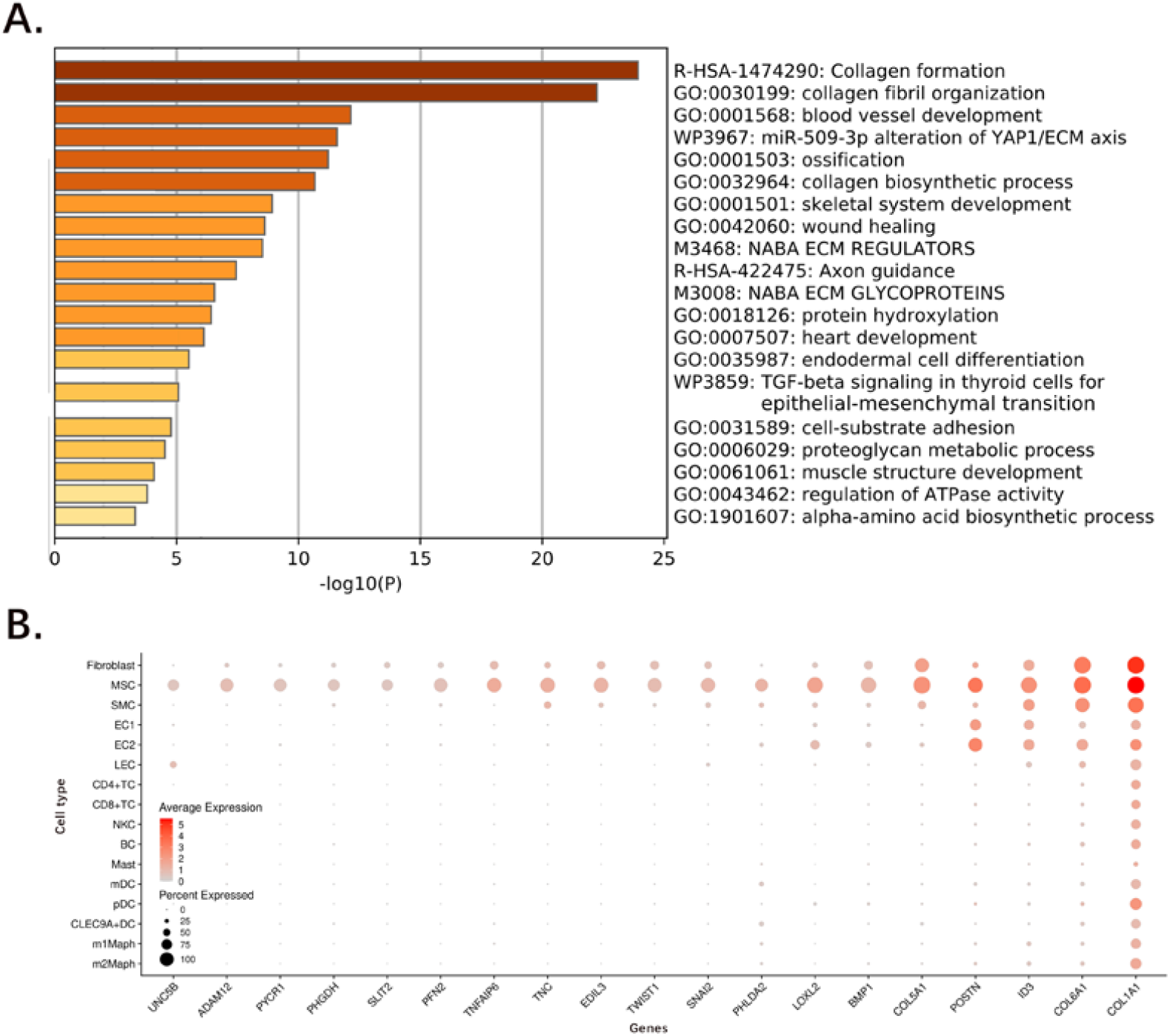
Identification of mesenchymal stem cells. The differential expression analysis between the cells inside and outside fibroblast2 was performed to generate a gene-expression signature, including 63 coding genes and a long noncoding RNA (lncRNA) gene RP11-14N7.2 (**Table 2**). **A**. The GO and pathway annotation of the 63 coding genes with analysis were performed using the Metascape website; **B**. Further investigation of the 63 coding genes (**Table 2**) showed at least 19 genes are over-expressed or up-regulated in stem cells. Among these 19 genes, *PYCR1, TNFAIP6, EDIL3, TWIST1, LOXL2, BMP1*, and *COL1A1* have been reported to be expressed in mesenchymal stem cells (MSCs) in the previous studies (**Supplementary file 3**).

**Table 2.**
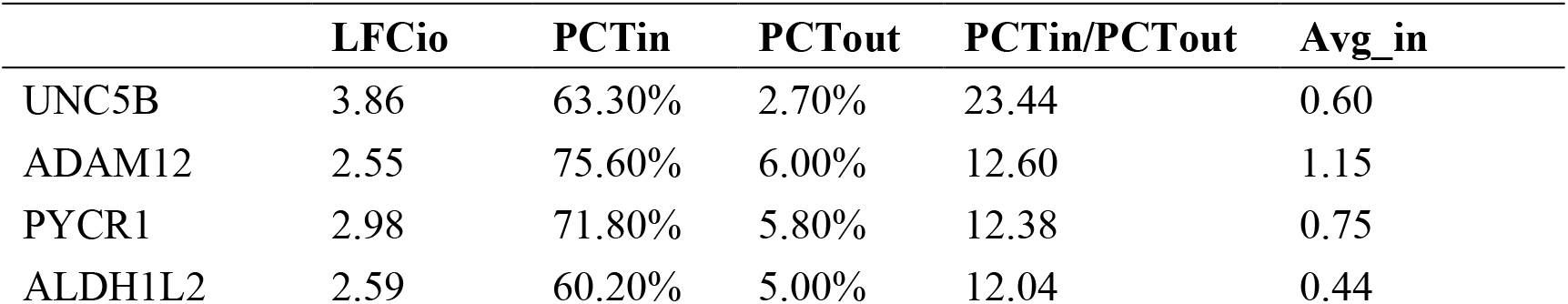
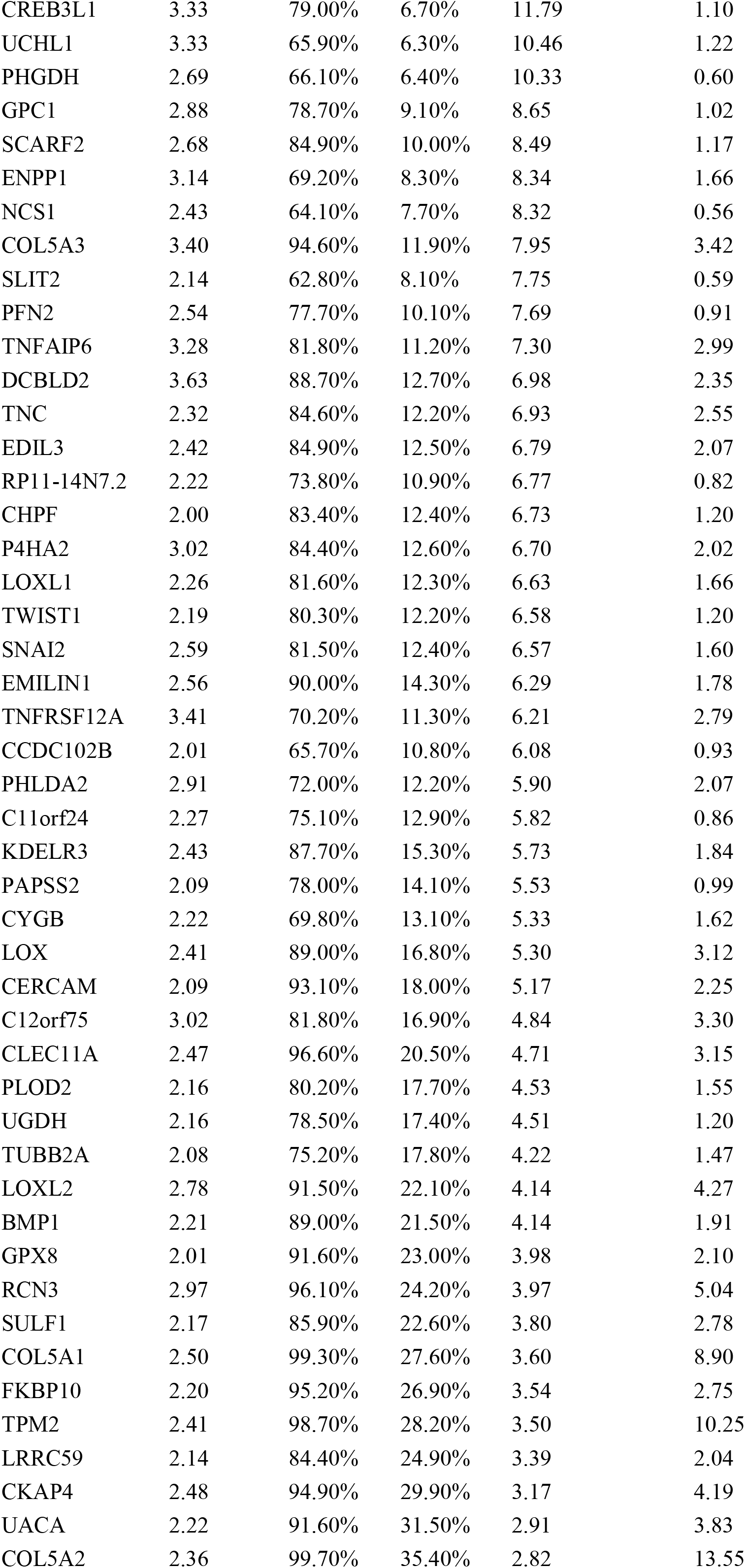
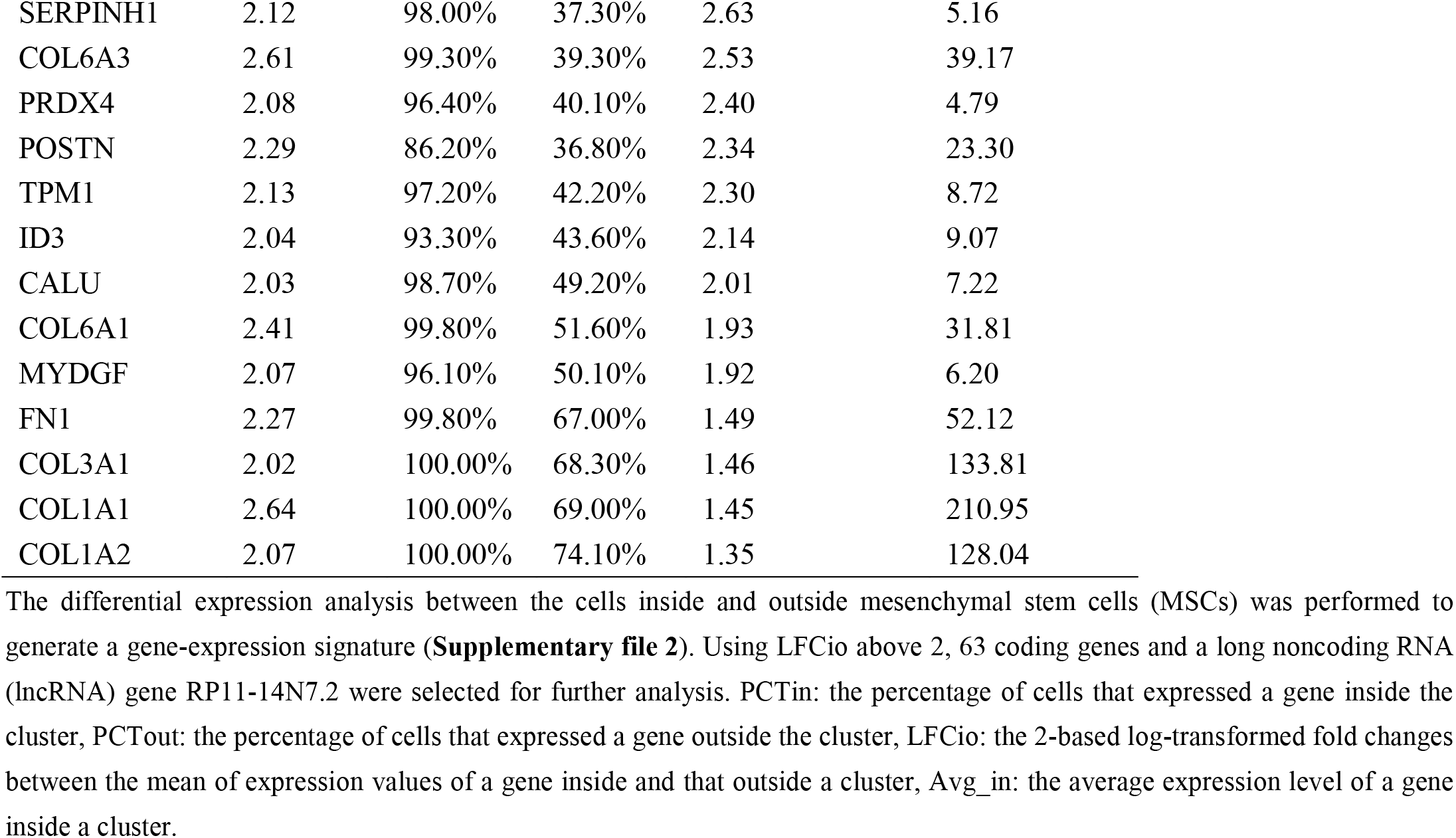
The gene-expression signature of mesenchymal stem cells.

Further investigation revealed that at least 33 genes in the gene-expression signature (**Table 2**) are currently being studied in cancers. Among these genes, 25 genes (*UNC5B, ADAM12, PYCR1, PHGDH, GPC1, ENPP1, NCS1, PFN2, TNFAIP6, DCBLD2, EDIL3, P4HA2, TWIST1, SNAI2, KDELR3, CERCAM, CLEC11A, PLOD2, UGDH, LOXL2, GPX8, SERPINH1, PRDX4, POSTN* and *MYDGF*) were over-expressed or up-regulated in cancers (**Figure S6**), whereas 7 genes (*CREB3L1, SLIT2, CHPF, TNFRSF12A, SULF1, UACA* and *TPM1*) were under-expressed or down-regulated (**Figure S7**). *UCHL1* acts as a tumour promoter in pancreatic, prostate, and lung cancers and as a tumour suppressor in ovarian cancer, hepatocellular cancer, and nasopharyngeal carcinoma. Herein, *UCHL1* was identified as the best marker gene for discriminating MSCs from other cells (**Figure S7**). The top highly expressed genes *COL1A1, COL3A1, COL1A2, FN1, COL6A3, COL6A1, POSTN, COL5A2, TPM2*, and *ID3* (**Figure S8**) were further analysed to derive new insights into tumour initiation, progression, and metastasis. Notably, *POSTN* was identified to be expressed at very high levels in MSCs, at high levels in the EC2 cluster, medium levels in the EC1 cluster, and low levels in fibroblasts. *POSTN* encodes a secreted extracellular matrix protein that functions in tissue development and regeneration, including wound healing and ventricular remodelling following myocardial infarction. According to a previous study [17], *POSTN* is expressed by fibroblasts in normal tissue and the stroma of the primary tumour and plays a role in cancer stem cell maintenance and metastasis. In humans, high expression of *POSTN* has been detected in various types of cancer, including breast, ovarian, lung, prostate, kidney, intestine, and pancreas [18]. Another previous study [19] reported *POSTN* over-expression in CAFs, which suggests that *POSTN* constitutes the primary tumour niche by supporting cancer cell proliferation through the ERK signalling pathway in gastric cancer. In the present study, *POSTN* was found to be expressed in MSC, EC2, and EC1 clusters at levels from highest to lowest (**described below**), which expands our understanding of the origin and functions of *POSTN* in stem cell differentiation, development, and cancer.

### Two clusters of endothelial cells

The EC1 and EC2 clusters contained 9.57% (1,032/10,784) and 21.08% (2,273/10,784) of the total cells, respectively. Among the top five marker genes of EC1 (**Table 1**), *TFF3* and *MMRN1* were highly expressed in LECs, whereas *EFNB2, RAPGEF5*, and *TIE1* were highly expressed in EC2 (**Figure 2B**). *MKI67* (marker of Ki-67 proliferation), *HGF, VEGFA, VEGFB*, and *VEGFC* were expressed at very low levels in the EC1 and EC2 clusters, whereas other *VEGF* and *FGF* genes (e.g., *FGF1, FGF2, FGF5, FGF9, FGF10, FGF11, FGF13, FGF14, FGF16, FGF18*, and *FGF22*) were barely detected in the EC1 and EC2 clusters (**Figure S9**). Such finding explains the phenomenon that “normal” vascular endothelial cells can be observed in the lesions of cavernous hemangiomas (**Introduction**). Unexpectedly, *VEGFA* and *VEGFB* were found to be expressed at very high levels in the m1Maph and pDC clusters, respectively. *MKI67*, a tumour proliferation marker, encodes Ki-67, which is associated with cellular proliferation (**Figure 3)**. Accordingly, the expression of *MKI67* correlates with tumour grade in many cancers. As cavernous hemangiomas are not malignant tumours, *MKI67* is not supposed to be detected in the lesion. However, *MKI67* was found to be expressed at low levels in the MSC cluster and was hardly detected in other cells. In addition, *GAPDH* expression was significantly higher in the MSC cluster than in the other cells. Such findings indicate that MSCs cause tumour formation in cavernous hemangiomas.

**Figure 3.**
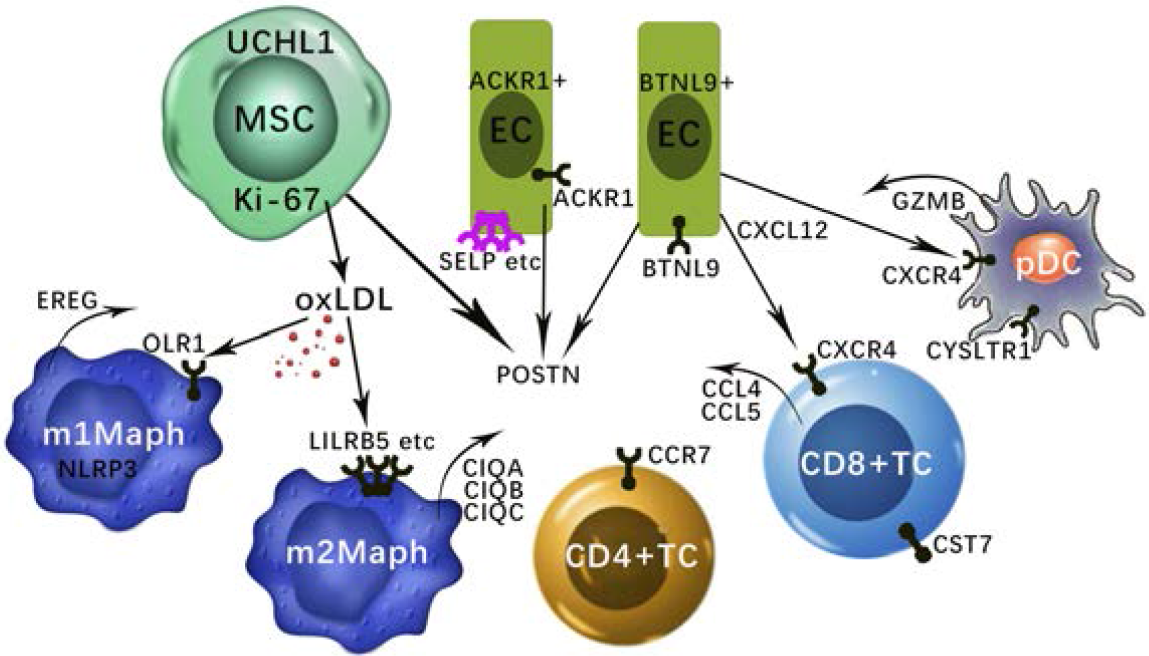
New discoveries in cavernous hemangioma. Mesenchymal stem cells (MSCs) that caused the tumor formation in the cavernous hemangioma. *UCHL1* is the best marker gene to discriminate these MSCs from other cells. *MKI67* (encoding Ki-67) was expressed at low level in the MSCs and hardly detected in other cells. MSCs induced the responses of BTNL9 positive endothelial cells (ACKR1+ECs) and BTNL9 positive endothelial cells (BTNL9+ECs). Adhesion molecules including SELP, SELE, and VCAM1 are up-regulated leading to leukocyte transmigration across the endothelium to the site of inflammation. *POSTN* was expressed at very high levels in MSCs, at high levels in the EC2 cluster, and at medium levels in the BTNL9+ECs and ACKR1+ECs. *CCR7* can be used as a better marker to discriminate the CD4+T cells from the activated CD8+T cells. CD8+T cells and NK cells may highly express *CCL5* for their infiltration in cavernous hemangiomas, independent on the tumor cell-derived *CCL5*-*IFNG*-*CXCL9* pathway. The highly expressed *BTNL9* in BTNL9+EC may cause checkpoint blockade and the effect was enhanced by the CXCL12-CXCR4 signalling. The highly co-expression of *CXCR4* and *GZMB* suggested that pDCs function for anti-tumour as CD8+T cells. We propose that oxLDL induces the oxLDL-OLR1-NLRP3 pathway by over-expression of *OLR1* in M1-like macrophages, whereas oxLDL induces the oxLDL-SRs-C1q pathway by over-expression of other scavenger receptors (*LILRB5*, etc) in M2-like macrophages.

Differential expression analysis (**Supplementary file 2**) between the EC1 and EC2 clusters revealed that almost all the major histocompatibility complex (MHC) class II genes (particularly, *HLA-F, HLA-DMA, HLA-DMB, HLA-DPA1, HLA-DPB1, HLA-DQA1, HLA-DQB1, HLA-DRA, HLA-DRB1*, and *HLA-DRB5*) were expressed at significantly higher levels in EC1 than in EC2 (**Figure S10**). As for other MHC II genes, *HLA-DOA, HLA-DOB, HLA-DQA2*, and *HLA-DQB2* were expressed at very low levels in the EC1 and EC2 clusters and *HLA-A, HLA-B, HLA-C*, and *HLA-E* were expressed at very high levels in all cell types except the mast cell. According to previous studies [20], MHC I genes are expressed by all nucleated cells, while the expression of MHC II genes is limited to antigen-presenting cells (APCs). Professional APCs (e.g., BCs, DCs, and macrophages) ubiquitously express MHC II, while cells such as ECs, which are not considered classic APCs, can induce MHC II expression in response to stimulation. The above results indicate that EC1 is a cluster of immune-responsive ECs. Various factors and stimuli, including cytokines, can induce type I activation of ECs, a state of heightened responsiveness. These responses are mediated by the binding of ligands to the extracellular domains of heterotrimeric G protein-coupled receptors (GPCRs) [20], such as *ADRA1D, ADRA2B, ADRA2C, AVPR2, CNR1, GALR1, GPR20, LPAR4*, OXTR, *P2RY2, TACR1, CYSLTR1, TAS1R1, TAS2R43, MAS1, GPR156, P2RY8, GPR52*, and *GPR85* (**Figure S11**). However, all of these GPCRs were expressed at very low levels or were barely detected in the EC1 and EC2 clusters. Various factors and stimuli can also induce type II activation of ECs, which is a relatively slower response that depends on new gene expression but delivers a more sustained inflammatory response [20]. The type II activation of ECs in cavernous hemangiomas should be further investigated in future studies.

EC1 was further clustered into two subtypes: *ACKR1* positive endothelial cells (ACKR1+EC) and *BTNL9* positive endothelial cell (BTNL9+EC), including 368 and 664 cells, respectively. *ACKR1, SELP, SELE, VCAM1*, and *CADM3* were expressed at significantly higher levels in ACKR1+EC than in BTNL9+EC (**Figure S12**). According to annotations from the GeneCards database [15], *ACKR1* encodes a glycosylated membrane protein as a non-specific receptor for several chemokines and may regulate chemokine bioavailability and consequently, leukocyte recruitment through two distinct mechanisms. As adhesion molecules, *SELP, SELE*, and *VCAM1* were up-regulated, leading to leukocyte transmigration across the endothelium to the site of inflammation [20]. However, another adhesion molecule *ICAM1* was not detected. *CADM3* encodes a calcium-independent cell-cell adhesion protein that can form homodimers or heterodimers with other nectin proteins. According to a previous study [20], these adhesion molecules can be induced by oxidised low-density lipoprotein (oxLDL). The expression of a scavenger receptor of oxLDL encoded by *OLR1* has been shown to be up-regulated in response to stimulation by oxLDL, pro-inflammatory cytokines, and proatherogenic factors such as angiotensin II in ECs. However, *OLR1* and genes encoding other scavenger receptors (e.g., *LILRB5, MRC1, MSR1, CD68, CD163, CXCL16*, and *CLEC7A*) were barely detected or (e.g., *STAB1* and *CD36*) expressed at very low levels in the EC1 and EC2 clusters (**Figure S13**).

*BTNL9* and *CXCL12* were expressed at significantly higher levels in BTNL9+ ECs than in ACKR1+ECs (**Figure S12**). As a member of the BTN/MOG Ig-superfamily, the protein encoded by *BTNL9* is expressed in a variety of tissues in humans and mice and functions as a negative regulator of immune cell activation. Recombinant BTNL9–Fc has been demonstrated to bind to many immune cells, including macrophages, T, B, and dendritic cells. In particular, *BTNL9* has been reported to inhibit CD8+ T-cell proliferation [21]. According to previous studies, *CXCL12* encodes a stromal cell-derived alpha chemokine member of the intercrine family and stimulates the migration of monocytes and T-lymphocytes through its receptors, *CXCR4* and *ACKR3* (**Figure S12**). In the present study, *CXCL12* was found to be highly expressed in the following non-immune cells in order of highest to lowest levels: fibroblast, MSC, SMC, EC1, EC2, and LEC clusters, while *CXCR4* was expressed at high levels in all immune cells, except mast cells. *ACKR3* was expressed at very low levels in all non-immune cells, but barely detected in immune cells. The above results suggest that the high expression of *BTNL9* in BTNL9+EC may cause checkpoint blockade and is enhanced by *CXCL12-CXCR4* signalling.

### T lymphocytes and NKCs

The TC1, TC2, and TC3 clusters that contained 5.53% (596/10,784), 2.85% (307/10,784), and 1.86% (201/10,784) of the total cells, respectively, were further identified as CD4+TC, activated CD8+TC, and NKC clusters. The cells in these three clusters were entangled, as they expressed some common genes. For example, *CD8B* (a marker gene of CD8+T) and *CD4* (a marker gene of CD4+T) were expressed at low levels in the CD4+T cells and the activated CD8+T cells, respectively. Further, both the activated CD8+T and the NK cells express *NKG7* at very high levels. Therefore, we compared the expression levels of more relevant genes to confirm the cell types of the three clusters. The NK cells were confirmed based on the following evidence: (1) the average expression levels of *CD3D, CD3E, CD3G, CD4, CD8A*, and *CD8B* (the marker genes of T lymphocytes) in the NK cells were lower than 5% of those in the CD4+T cells and activated CD8+T cells, respectively (**Figure S14**); (2) the average expression level of *NKG7* in the NK cells was approximately 92-fold higher than that in the CD4+T cells; and (3) the average expression levels of *GZMA, GZMB, GZMH, GZMK, GZMM, CRTAM*, and *GNLY* in the NK cells were approximately 12.3, 51, 4.56, 102.2, 1, 11.2, and 151-fold higher than those in the CD4+T cells, respectively (**Figure S14**). Although both the CD4+T cells and activated CD8+T cells expressed *CD8B* at a similar level, the CD4+T cells could still be distinguished from the activated CD8+T cells based on the following evidence: (1) the average expression level of *CD4* in the CD4+T cells was higher than 5-fold of that in the activated CD8+T cells; (2) the average expression level of *CD8A* in the activated CD8+T cells was higher than 4-fold of that in the CD4+T cells; (3) the average expression levels of *NKG7, GZMA, GZMB, GZMH, GZMK, GZMM, CRTAM*, and *GNLY* in the CD4+T cells were markedly lower than those in the NK and activated CD8+T cells; and (4) the average expression level of *CCR7* in the CD4+T cells was higher than 38-fold of that in the activated CD8+T cells and 20-fold of that in the NK cells. Herein, we revealed for the first time that: (1) *CCR7* is expressed at a markedly higher level than *CD4, CD8A*, and *CD8B* and can be a better marker to distinguish CD4+T cells from activated CD8+T cells; and (2) the average expression levels of *GZMA, GZMH, GZMK*, and *GZMM* in the activated CD8+T cells are higher than 2-fold of those in CD4+T cells, while the average expression levels of *GZMB, CRTAM*, and *GNLY* are only approximately 53.1%, 35.7%, and 15% of those in the NK cells, respectively.

Differential expression analysis (**Materials and Methods**) between cells inside and outside the activated CD8+T cluster revealed 44 coding genes as a gene-expression signature (**Supplementary file 3**), which merits further investigation. Among the top five highly expressed genes, *B2M, CCL4, RPS27, RPS15A*, and *CCL5* (**Figure S15**), both *CCL4* and *CCL5* encode secreted chemokine ligands that have chemokinetic and inflammatory functions by binding to their receptor encoded by *CCR5. CCL4* can promote tumour development and progression by recruiting regulatory T cells and pro-tumorigenic macrophages, and acting on other resident cells (e.g., fibroblasts and endothelial cells) present in the tumour microenvironment (TME) to facilitate their pro-tumorigenic capacities [22]. In contrast, in some situations, *CCL4* can enhance tumour immunity by recruiting cytolytic lymphocytes and macrophages with phagocytic ability. The over-expression of *CCL5* is associated with CD8+T cell infiltration in solid tumours [23]. T cell infiltration requires tumour cell-derived *CCL5*, and this process is amplified by IFN-γ-inducible, myeloid cell-secreted *CXCL9*. As only IFN-γ encoded by *IFNG* was detected in ovarian cancers [23], we named this amplification process as tumor cell-derived *CCL5-IFNG*-*CXCL9* pathway. By examining the expression levels of *CCL4, CCL5, CCR5*, and *CXCL9* in all immune and MSCs (**Figure S15**), we found that: (1) *CCL4* was expressed at very high levels in the CD8+TC cluster and at high levels in the NKC, m1Maph, and 2Maph clusters; (2) *CCL5* was expressed at very high levels in the CD8+TC cluster and at medium levels in the NKC cluster; and (3) *CCR5, IFNG*, and *CXCL9* were expressed at very low levels or not detected in the present study. These results suggest that CD8+T cells and NK cells highly express *CCL5* for their infiltration in cavernous hemangiomas, independent of the tumour cell-derived *CCL5-IFNG*-*CXCL9* pathway. *CCL5* and *CXCL9* co-expression revealed immunoreactive tumours with prolonged survival and response to PD-1 inhibition [23]. *PDCD1* (well-known as *PD-1*) was barely detected in CD4+T cells and activated CD8+T cells. However, whether *PDCD1* functionally associate with *CCL5* is still unknown. *CST7* was also found to be expressed at high levels in activated CD8+T cells, rather than in tumour cells. However, the expression of *CST7* has been observed in various human cancer cell lines established from malignant tumours. According to annotations from the GeneCards database [15], *CST7* encodes a glycosylated cysteine protease inhibitor with a putative role in immune regulation through the inhibition of a unique target in the hematopoietic system. The specific functions of *CST7* in activated CD8+T cells merit further investigation.

### Two subsets of macrophages

A number of previous studies have revealed that a considerable degree of monocyte–macrophage heterogeneity exists when various marker genes are used to identify macrophage subsets [24]. An over-simplified generalisation of this concept recognises M1 and M2 macrophages, which play an important role in tumour progression. According to previous studies [25], M1 macrophages play a protective role, while M2 macrophages promote tumour growth. M1 macrophages mainly secrete proteins encoded by interleukin 12A (*IL12A*), interleukin 12 B (*IL12B*), and tumour necrosis factor (*TNF*), whereas M2 macrophages typically produce proteins encoded by interleukin 10 (*IL10*), interleukin 1 receptor antagonist (*IL1RN*), and interleukin 1 receptor type 2 (*IL1R2*) [26]. Other published marker genes for M1 macrophages include *IL1B* and *NFKB1*, while those for M2 macrophages include *MERTK, MRC1, STAB1*, and *CD163*. Using *IL1B, MERTK, MRC1, STAB1*, and *CD163* (**Figure S13**), two clusters of macrophages were barely identified and temporarily called M1-like and M2-like macrophages, respectively. However, no single marker gene can be used to discriminate M1-like macrophages from M2-like macrophages. For example, 65.84% (607/922) of M1-like macrophages and 88.18% (1134/1286) of M2-like macrophages highly express *CD163*. Therefore, a combination of marker genes (**Figure 1B**) must be used to confirm M1-like and M2-like macrophages.

In M1-like macrophages, 66.3%, 83.4%, 64.5%, 82.5%, and 71.3% of the cells expressed *OLR1, EREG, BCL2A1, SLC11A1*, and *NLRP3*, respectively (**Figure 1B**). A previous study reported that oxLDL-induced NLRP3 inflammasome activation (noted as the oxLDL-NLRP3 pathway) in macrophages plays a vital role in atherogenesis [27]. In the present study, the discovery of highly expressed *OLR1* and *NLRP3* revealed that oxidised LDL (oxLDL) induced NLRP3 inflammasome activation in M1-like macrophages through *OLR1* (noted as the oxLDL-OLR1*-*NLRP3 pathway). According to annotations from the GeneCards database [15], *OLR1*, a scavenger receptor of oxLDL, also mediates the recognition, internalisation, and degradation of oxLDL by vascular endothelial cells. Other highly expressed scavenger receptors of oxLDL [28], particularly those encoded by *MRC1, STAB1, MSR1, CD36, CD68, CD163, CXCL16*, and *CLEC7A* may not be involved in or may contribute minimally to the oxLDL-NLRP3 pathway, as they were also highly expressed with low expression of NLRP3 in M2-like macrophages. According to annotations from the GeneCards database [15], the protein, epiregulin, encoded by *EREG* is a ligand of the epidermal growth factor receptor (*EGFR*) and structurally related erb-b2 receptor tyrosine kinase 4 (*ERBB4*). *EREG* may be involved in a wide range of biological processes, including inflammation, wound healing, oocyte maturation, and cell proliferation. In particular, *EREG* promotes cancer progression in various human tissues. By single-cell transcriptome analysis, a previous study [29] revealed that *EREG* was predominantly expressed in macrophages in the TME and induced EGFR-tyrosine kinase inhibitor (TKI) resistance in the treatment of non-small cell lung cancer (NSCLC) by preventing apoptosis through the EGFR/ErbB2 heterodimer. In the present study, we found that M1-like macrophages highly express *EREG*, which provides a deeper understanding of the origin and functions of epiregulin in the TME.

In M2-like macrophages, 66.3%, 72.6%, 83.3%, 78%, and 85% of the cells expressed *FOLR2, LILRB5, C1QC, MS4A4A*, and *C1QB*, respectively (**Figure 1B**). The serum complement subcomponent, C1q, is composed of 18 polypeptide chains which include six A-chains, six B-chains, and six C-chains, encoded by complement C1q A chain (*C1QA*), complement C1q A chain (*C1QB*), and complement C1q A chain (*C1QC*) genes. The C1q protein enhances the survival and efferocytosis of macrophage foam cells [30], which is thought to be induced by low-density lipoproteins (LDL), including oxidised LDL (OxLDL) or minimally modified LDL (mmLDL). Understanding the molecular mechanisms involved in OxLDL- and mmLDL-induced macrophage foam cell formation is of fundamental importance for atherosclerosis and cardiovascular disease. However, whether M2-like macrophages include macrophage foam cells is still unknown. Another previous study [31] showed that the expression levels of *C1QA, C1Q*B, and *C1QC* were positively related to M1 and M2 macrophages and CD8+ cells, and negatively correlated with M0 macrophages. However, our results showed that *C1QA, C1QB*, and *C1QC* were expressed in M2-like macrophages at very high levels and M1-like macrophages at very low levels. The discovery of high expression levels of *LILRB5, C1QA, C1QB*, and *C1QC* suggested that oxLDL induced the inflammatory activation of C1q (noted as the oxLDL-SRs-C1q pathway, with SRs as genes encoding scavenger receptors of oxLDL, except *OLR1*) in M2-like macrophages via scavenger receptors *LILRB5, MRC1, STAB1, MSR1, CD68*, and *CD163*. Other scavenger receptors *CD36, CXCL16*, and *CLEC7A* may not be involved in or contribute slightly to the oxLDL-SRs-C1q pathway, as they were also highly expressed in M1-like macrophages; however, the oxLDL-SRs-C1q pathway is not induced. Notably, the oxLDL-OLR1-NLRP3 pathway is not induced in M2-like macrophages. Based on the above results, we propose that oxLDL induces the oxLDL-OLR1-NLRP3 pathway via the over-expression of *OLR1* in M1-like macrophages, whereas oxLDL induces the oxLDL-SRs-C1q pathway via the over-expression of other scavenger receptors in M2-like macrophages.

### Other cells

For the other five clusters, 0.35% (38/10784), 1.1% (119/10784), 2.47% (266/10784), 0.33% (36/10784), and 0.44% (47/10784) of the total cells were identified as LECs, BCs, mDCs, pDCs, and CLEC9A+DCs, respectively. Differential expression analysis (**Materials and Methods**) between cells inside and outside each cluster was performed to generate DE genes (**Supplementary file 2**) for further analysis. The top five DE genes were selected as the combination of marker genes for each cell type (**Figure 1B**). Of note, *CXCR4, GZMB*, and *CYSLTR1* were found to be expressed in the pDC cluster at very high levels (**Figure 3**). The high co-expression of *CXCR4* and *GZMB* suggests that pDCs function for anti-tumour as CD8+T cells in cavernous hemangiomas. Although the proportion of pDCs is markedly lower than that of CD8+T cells, their contribution to anti-tumour activity may complement the loss by checkpoint blockade in CD8+T cells. *CYSLTR1* encodes a protein that is a second receptor for cysteinyl leukotrienes and is thought to be the main receptor mediating cysteinyl leukotriene receptor smooth-muscle contraction and inflammatory cell cytokine production in asthma. However, the specific functions of the highly expressed *CYSLTR1* in pDCs remain unknown.

## Conclusion and Discussion

In the present study, we identified the 16 main cell types in cavernous hemangioma using scRNA-seq and assigned a combination of five top marker genes to each cell type. Such finding facilitates the repeated identification of these cell types in future studies. The main contribution of this study is the discovery of mesenchymal stem cells (MSCs) that cause tumour formation in cavernous hemangioma and we propose that these MSCs may be abnormally differentiated or incompletely differentiated from epiblast stem cells. We ruled out the possibility that the MSCs originated from pericytes or bone marrow-derived MSCs during the wound healing process, as the patient carried this tumour at birth and the MSCs did not express *NKD2, GREM2, NRP3*, and *FRZB*, which are markers of pericyte-to-myofibroblast differentiation.

MSCs in the cavernous hemangioma exhibit characteristics of tumors, including the over-expression of *MKI67* and *GAPDH* and the oxLDL induced pathways. *UCHL1* is the best marker gene to discriminate these MSCs from other cells. MSCs induced the responses of ECs, although they are usually observed as “normal” in cavernous hemangiomas by microscopic examination. Two immune-responsive ECs were identified as ACKR1+EC and BTNL9+EC. The highly expressed *BTNL9* in BTNL9+EC may cause checkpoint blockade, and this effect was enhanced by *CXCL12-CXCR4* signalling. *POSTN* was expressed at very high levels in MSCs, high levels in the EC2 cluster, medium levels in the EC1 cluster, and low levels in fibroblasts.

The tumour cell-derived *CCL5-IFNG*-*CXCL9* pathway was not detected in cavernous hemangioma. CD8+T cells and NK cells may highly express *CCL5* for their infiltration into cavernous hemangiomas, independent of the tumour cell-derived *CCL5-IFNG*-*CXCL9* pathway. The specific functions of *CST7* in activated CD8+T cells warrant further investigation. *CCR7* is a better marker to distinguish CD4+T cells from activated CD8+T cells. The high co-expression of *CXCR4* and *GZMB* suggests that pDCs function for anti-tumour as CD8+T cells in cavernous hemangiomas. However, the specific functions of highly expressed *CYSLTR1* in pDCs remain unknown. M1-like macrophages highly express *EREG*, which provides a deeper understanding of the origin and functions of epiregulin in the TME. The oxLDL in the TME of cavernous hemangiomas may play an important role as a signalling molecular in the immune responses. Notably, we propose that oxLDL induces the oxLDL-OLR1-NLRP3 pathway by over-expression of *OLR1* in M1-like macrophages, whereas oxLDL induces the oxLDL-SRs-C1q pathway by over-expression of other scavenger receptors in M2-like macrophages.

## Materials and Methods

### 10x Genomics library preparation and sequencing

The tissues were digested for 0.5 h at 37 temperature in the enzyme solution (Enzyme H, R and A) using a gentleMACS Dissociator following the manufacturer’s instruction. The single-cell suspensions were filtered with a 40-μm-diameter cell strainer (FALCON, USA), then washed twice times with RPMI1640 wash buffer at 4 temperature. Cell viability was determined by trypan blue staining with TC20 automated cell counter (BioRad, USA). The ratio of viable cells in single-cell suspension was more than 85%. The concentration of single-cell suspension was adjusted to 700–1200 cells/μL. The cells of each sample were then processed to the generate 10x libraries with the Chromium Single Cell 3’ Reagent Kits v3 CG000183 (10x Genomics, USA) following the manufacturer’s instruction. These libraries were sequenced by a Illumina NovaSeq sequencer, producing 542,088,397 pairs of 150-bp reads (∼ 163 Gbp raw data).

### ScRNA-seq data processing

Using Cell Ranger v4.0.0, we aligned 542,088,397 read2 sequences (∼) to the human genome GRCh38, generating an UMI-count matrix (33,694 genes × 6,794,880 cell barcodes). The cell-calling algorithm in Cell Ranger was used to identify 12,018 cells from the 6,794,880 cell barcodes. Then, a total of 10,784 cells and 22,021 genes were retained with a median of 2,084 genes per cell after quality control (QC) filtering using the following parameters: (1) genes detected in < 3 cells were excluded; (2) cells with < 200 genes were excluded; (3) cells with >30% mitochondrial RNA UMI counts or >5% hemoglobin RNA UMI counts were excluded; (4) 982 doublet artifacts were removed with DoubletFinder. Finally, a 22,010×10,784 matrix and a 11×10,784 matrix were produced to represent expression levels of nuclear and mitochondrial genes, respectively.

Seurat v4.0.1 was used for single cell data analysis on R v4.0.1 with other Bioconductor packages [32]. The nuclear UMI-count matrix were normalized per cell using the NormalizeData function by dividing the total number of reads in that cell, then multiplying by a scale factor of 10000 and taking natural-log transformed values. We selected 2000 highly variable genes on the basis of the average expression and dispersion per gene using the FindVariableFeatures function with parameters. After data scaling, the principal component analysis was performed on the 2000 highly variable genes using the RunPCA function. The top 50 principal components were chosen for cell clustering using the FindClusters function with resolution=0.4. The Uniform Manifold Approximation and Projection (UMAP) method was used to show the clustering results.

### Identification of cell types and selection of marker genes

Several differentially expressed (DE) genes were selected for each cluster, satisfying the following criteria: (1) the adjusted p-value < 0.01; (2) the percentage of cells that expressed the gene inside the cluster (PCTin) > 60%; and (3) the ratio between the percentage of cells that expressed the gene inside (PCTin) and that outside the cluster (PCTout) is ranked in the top five. LFCio is the 2-based log-transformed fold changes between the mean of expression values of a gene inside and that outside a cluster. The differential expression analysis between cells inside and outside the cluster was performed using the R package DESeq2. To use DESeq2 for the scRNA-seq data analysis, we calculated the size factors using scran::computeSumFactors with the following parameters: test = “LRT”, fitType = “glmGamPoi”, minReplicatesForReplace = Inf, useT = TRUE, minmu = 1e-6. Genes with expression levels below 10% in both of the two groups of cells, were filtered out. The GO and pathway annotation with analysis were performed using the Metascape website (https://metascape.org/gp) [33].

For each cluster, we identified its cell type by comparing the selected DE genes to known marker genes. As for the known marker genes, (1) most of them were used according to records in a online database (http://biocc.hrbmu.edu.cn/CellMarker/) by Harbin Medical University and (2) a few were used according to records in published papers, e.g. LINC00926 [34]. As LFCIO and PCTDIO can only be used to evaluate the identification of cell type by a single marker gene. we designed another metric UICC to evaluate the representation of a cell type by a combination of marker genes. The cardinal number of the union set of cells expressed marker genes is divided by the number of all cells in a cluster to calculate the union coverage of a cluster (UCC). The cardinal number of the intersection set of cells expressed marker genes is divided by the number of all cells in a cluster to calculate the intersection coverage of a cluster (ICC). To balance the UCC and ICC, we designed UICC, which is calculated by multiplying UCC by ICC.

## Supplementary information

### Declarations

#### Ethics approval and consent to participate

Not applicable.

#### Consent to publish

Not applicable.

#### Availability of data and materials

All data used in the present study was download from the public data sources.

#### Authors’ contributions

Shan Gao conceived the project. Shan Gao and Daqing Sun supervised this study. Guangyou Duan and Jia Chang performed programming. Xin Li and Qiang Zhao, Jinlong Bei and Tung On Yau downloaded, managed and processed the data. Jianyi Yang predicted the protein structures. Shan Gao drafted the main manuscript text. Shan Gao and Jishou Ruan revised the manuscript.

## Acknowledgments

We are grateful for the help from the following doctors at Tianjin Medical University General Hospital: Wenjing Song, Xiaohui Liang from from the pathology department; Junping Wang from the medical imaging department. We would like to thank Editage (www.editage.cn) for polishing part of this manuscript in English language. This manuscript was online as a preprint on xx xx, 2021 at yy.

